# Enhancing drug repurposing on graphs by integrating drug molecular structure as feature

**DOI:** 10.1101/2023.05.03.539227

**Authors:** Adrián Ayuso-Muñoz, Lucía Prieto-Santamaría, Andrea Álverez-Pérez, Belén Otero-Carrasco, Emilio Serrano, Alejandro Rodríguez-González

## Abstract

Drug repurposing has become increasingly important, particularly in light of the COVID-19 pandemic. This process involves identifying new therapeutic uses for existing drugs, which can significantly reduce the cost, risk, and time associated with developing new drugs, *de novo* development. A previous conducted study proved that Deep Learning can be used to streamline this process by identifying drug repurposing hypotheses. The study presented a model called REDIRECTION, which utilized the rich biomedical information available in graph form and combined it with Geometric Deep Learning to find new indications for existing drugs. The reported metrics for this model were 0.87 for AUROC and 0.83 for AUPRC. In this current study, the importance of node features in GNNs is explored. Specifically, the study used GNNs to embed two-dimensional drug molecular structures and obtain corresponding features. These features were incorporated into the drug repurposing graph, along with some other enhancements, resulting in an improved model called DMSR. Performance score for the reported metrics values raised by 0.0448 in AUROC and 0.0919 in AUPRC. Based on these findings, we believe that the method used for embedding drug molecular structures is interesting and captures valuable information about drugs. Its incorporation in the graph for drug repurposing can significantly benefit the process, leading to improved performance evaluation metrics.

## I. Introduction

In our prior work, Ayuso *et al*. (2022) [1], it was proved that incorporating diverse biomedical data into a multi-layered graph is a helpful method for gaining a deeper understanding of diseases and their associated components and characteristics. Using Graph Neural Networks (GNNs) and DISNET biomedical graph [2] a model called REDIRECTION was presented. DISNET is a biomedical integrated knowledge base containing information regarding diseases and their associations to symptoms and drugs, among others.

REDIRECTION is based on graph deep learning [3], [4] and link prediction. Its aim is to generate drug repurposing hypotheses. Drug repurposing or repositioning involves identifying new therapeutic uses for drugs that have already been approved. The model performed well, scoring 0.87 for the area under the receiving operating characteristics curve (AUROC) and 0.83 for the area under the precision-recall curve (AUPR) for a selected subset of already-tested repurposing hypotheses, RepoDB test.

GNNs benefit enormously from the addition of node features, due to the foundational role these features play in the message passing framework upon which GNNs are built [5], [6]. This is an intuitive outcome, as the incorporation of relevant information into the dataset is likely to enhance the model’;s performance.

Hence, in the context of GNNs, the two-dimensional molecular structure of the drugs in the dataset will be embedded into a vector, which will be used as node feature of the corresponding drug on the drug repurposing hypotheses generation. The drug molecular structure, which is represented as a graph structure depicting the arrangement of atoms and bonds in a molecule of a drug, thus plays a pivotal role in this process.

The main contribution introduced in current research is the refinement of the techniques used to develop the model and the verification that node features play a fundamental role in the GNNs paradigm. These advancements ultimately result in an improved model called DMSR (Drug Molecular Structure Redirection).

The paper is organized as follows: Section II includes a brief revision of how GDL has been previously employed in the context of drug molecular structure embedding, Section III details how the heterogeneous graph has been built, how the drug molecular structure has been generated, the specifications of the models and how they have been evaluated and validated. Section IV depicts the different results obtained, while Section V finishes with the conclusions and future lines.

## II. Related work

Molecular representation has been a topic of interest since the nineteenth century [7]. In the late years, with the boom of deep learning, this field has undergone a significant transformation [8]–[12]. Specifically, GNNs [13], [14] are particularly well-suited for molecular representation due to their ability to handle graphs directly. In fact, a comparison of different molecular machine learning models conducted in Wu *et al*. (2018) [14] demonstrated that GNNs outperform other approaches by a considerable margin on most datasets. One interesting observation is that, while some studies focus on proteins rather than drugs [15] or on both [16], the underlying problem is essentially the same.

To the best of our knowledge, the first paper to use the message passing framework for molecular representation is Gilmer *et al*. (2017) [13]. This paper introduces a framework for supervised learning on molecules, demonstrating the ability of message passing neural networks to predict molecular properties and eliminating the need for complex feature engineering.

The objective is using this drug molecular representation as a steppingstone to accomplish a task. Various studies have utilized this representation for different purposes, such as drug discovery [7], [17], molecule design [18], [19] or drug repurposing [20].

Sanchez-Lengeling and Aspuru-Guzik (2018) [18] conducted a review of methods for molecular design starting from a set of desired properties. The authors analyzed various molecular generative models, including variational autoencoders, generative adversarial networks, reinforcement learning, recurrent neural networks, and hybrid approaches, to represent the latent functional space.

Similarly, in Seo *et al*. (2023) [19], a graph-based generative model was introduced which constructs molecules by concatenating prepared chemical building blocks until the target properties are achieved. GNNs were employed to compute certain properties of a given molecular structure.

Aligned with the aim of this work, in Pham *et al*. (2021) [20], DeepCE was presented. This model uses simplified molecular-input line-entry system (SMILES) to create a graph and then applies a Graph Convolutional Network (GCN) to generate features for the drug. The model then sums up all the node representations in the graph to produce the final vector. The goal is to generate hypotheses for drug repurposing by utilizing the features of the chemical compounds, along with information related to genes, cell-specific data, and drug dosage.

## III. Materials and methods

In this section, we describe how the input graphs for our models are generated, as well as the model architecture, training, testing, and validation procedures. We also discuss our secondary validation process.

### A. Materials

#### 1) Drug Repurposing Graph

The used graph for the drug repurposing task is represented in Fig. 1. It is constructed the same way as the previous version was, using DISNET database data. The graph consists of 30,731 *phenotype* nodes and 3,944 *drug* nodes, with 52,179 *dis_dru_the* and 313,972 *dse_sym* edges. For further information, please refer to Ayuso *et al*. (2022) [1]. Notably, the edges are now undirected.

**Fig. 1.**
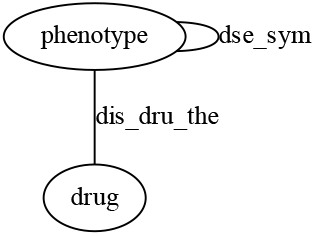
Structure of the heterogeneous graph.

The calculated vectors by the drug molecular structure embedding model will be added as features for the drug nodes, having a vector of ones in case of absence of SMILES representation. To assess the impact of these features on performance, two datasets will be created - one with the added features and another without - for a comparative analysis.

It is notable that a portion of the *dis_dru_the* edges are removed from the graph before splitting it to train, test and validate, to avoid data leakage. This edges correspond to the information present in RepoDB [21], which contains a standard set of drug repositioning successes and failures that can be used to benchmark computational repositioning methods.

#### 2) Drug Molecular Structure

The molecular representation of all drugs in the DISNET database is obtained by transforming their SMILES representation to two-dimensional graphs through the use of the *pysmiles* library [22]. This returns molecules as a *NetworkX* [23] graph, where each node has the following features: element, charge, aromatic, hcount (implicit hydrogens attached) and stereo (stereochemical information). For drugs that do not have SMILES representation in the DISNET database, the SMILES representation is retrieved from PubChem [24] through the use of *PubChemPy* [25]. Following this step, 364 drugs still lack a SMILES representation and are thus excluded from the training dataset, having 3,580 molecular structures.

SMILES is a line-based notation system for chemical structures, based on graph theory and using a predefined grammar [28].

### B. Methods

All the development was carried out in Python 3.8.10, under the framework of the Stanford library *DeepSNAP 2*.*0* (https://snap.stanford.edu/deepsnap/), *PyTorch Geometric 2*.*0*.*3* and *NetworkX 2*.*6*.*3*. We made use of CUDA Toolkit 11.3, running the experiments on an Ubuntu Server LTS 20.04.4 with a GPU (NVIDIA GeForce RTX 3090 24GB), Intel i9-12900K and 32GB RAM. All code and results have been published in an accessible repository^1^.

#### 1) Drug Repurposing Model

The drug repurposing model architecture follows the same pipeline as the one presented in Ayuso *et al*. (2022) [1]. Its training, testing, validating and RepoDB phase have not changed. It uses GraphSAGE layers to generate embeddings, encoder, and the sigmoid function applied over the dot product of two embeddings as a decoder. The hyperparameters are kept the same for the three models so the comparison is fairer: 200 epochs, 1e^-4^ weight decay, 0.01 learning rate, 0.8 edge message passing rate and 32 dimensions for embeddings.

The model is comprised of a two-layer GNN that encodes the nodes in the biomedical graph. It then calculates the dot product between pairs of drugs being considered for repurposing. The sigmoid of the dot product output represents the probability of an edge existing between the pair, indicating the therapeutic potential of a drug for a specific disease.

#### 2) Drug Molecular Structure Model

The drug molecular structure model employs an unsupervised learning autoencoder architecture, specifically self-supervised. The applied model is based on the one in Kipf and Welling (2016) [26]. Since there are no assigned classes or labels for the molecular structures, the goal is to compress them into a compact vector that can serve as features, called code. Once it has been trained, just the encoder section will be kept since the code is the useful part. A schema of the model’ information flow: first the drug input, then SMILES representation, following the graph representation and finally the encoded drug, can be seen in Fig. 2.

**Fig. 2.**
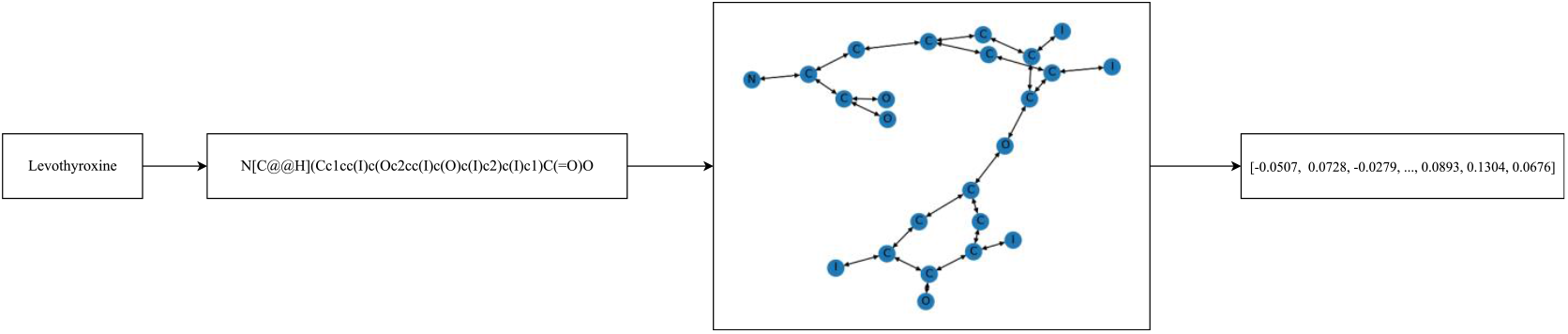
Schematic representation of the information flow in the Drug Molecular Structure model. Having a drug as an input, its SMILES code is retrieved and then transformed into a graph. That graph is embedded using GNNs into a vector that represents the whole molecular structure. The letters seen in the graph correspond to the chemical element of that node. In this image, Levothyroxine is used as an example.

Autoencoders are divided into two parts: i) encoder, which compresses the input into a code, and ii) decoder, which decompresses the code trying to match the original input. In this work, the autoencoder architecture is used for dimensionality reduction and data compression. Other applications for autoencoders include noise reduction or anomaly detection.

The autoencoder’s encoder consists of two GraphSAGE layers for encoding the graph, with a leaky rectified linear unit (LRELU) in between. The input is the graph representing the molecular structure of the drug, with a 5-dimensional feature vector per node, as it is the information the library can extract from the SMILES representation, described previously, and the code is a 32-dimensional vector.

The decoder is automatically generated by the interface graph autoencoder (GAE) of *PyTorch Geometric* [27]. It applies a sigmoid function to the inner product between the code and its transpose, that is, the probability of the existence of an edge between node-pairs. It is only used during the training phase.

During training, the loss function computes the binary cross entropy with logits loss (BCELogitsLoss) for both positive and negative edges. The model is trained along 200 epochs, using 1e^-3^ as learning rate, 20% of the molecules are reserved as the test set, without a validation split.

Once it has been trained and tested, all the dataset molecules are encoded using the model and saved in a file to be used as features in the drug repurposing task. Molecules that do not have a SMILES code have a 32-dimensional vector full of ones. The graph embeddings are generated by calculating the mean of all the node’s embedding.

It is important to note that autoencoders, as an inherent characteristic, may not function accurately for data that deviates significantly from the distribution used during training, resulting in unpredictable outcomes.

#### 3) Evaluating and validating hypotheses

The models were assessed using a separate test set during the training phase to verify the correctness of the training process. The training set comprises 80% of the original data and is distinct from the test data, which accounts for 20% in the drug molecular embedding model and 10% in the other model. Negative edges are randomly selected, which may cause some issues in the drug repurposing model if there are potential positive edges between the sampled ones.

Initially, the models were assessed using traditional performance metrics for the test set, such as the area under the receiver operating characteristic curve (AUROC) and the area under the precision-recall curve (AUPRC), as well as their graphical representation. AUROC measures the balance between true positive and false positive rates at various threshold levels, while AUPRC is computed based on precision and recall values at different threshold levels.

Secondly, the drug repurposing model underwent an additional validation by performing real drug repurposing cases. These cases were taken from RepoDB, and cases already present in the DISNET database were removed to prevent data leakage. A total of 5,013 repurposing cases were sampled to conduct the secondary validation, using the same metrics as the previous phase to evaluate the model’;s performance. Additionally, 5,013 drug-disease pairs were randomly sampled to ensure that the model did not assign high scores arbitrarily.

In addition, an analysis was conducted on the distribution of scores for both RepoDB and random links. It was expected that the scores for edges in the random set would be close to 0, while those in the RepoDB set would be closer to 1.

## IV. Results and discussion

The presented results for the drug repurposing model are obtained using multiple resampling-based estimation methods during training time and reporting the results obtained on the RepoDB test, so there is certain stability of the estimates. Values are estimated through repeated hold-out with k=100.

The hold-out estimation approach involves dividing a dataset into separate and non-overlapping sections, typically training and testing and sometimes validation (for various development tasks). The algorithm’;s parameters are adjusted based on the training set, and the model is then evaluated using the test set to obtain performance metrics. To improve the accuracy of the estimates, repeated hold-out technique is utilized, which involves repeating the process multiple times with random assignment of the sets each time. This reduces the estimates’; variability but does not offer control over how frequently a sample is assigned to either set.

Confidence intervals (95%) are calculated using the normal distribution, since the number of samples is over 30, there are 100 samples. REDIRECTION model has no confidence interval since it was presented as an unique model and there is no other available data.

TABLE I shows a comparison of the models’; results. The latest model, DMSR, significantly improves the previous version by considering truly undirected edges, allowing better adjustment and node representations due to bidirectional information flow. Adding features to DMSR -WF (without features) leads to even greater improvement, with an increase of 0.0448 in AUROC and 0.0919 in AUPRC compared to REDIRECTION, and an increase of 0.012 in AUROC and 0.0042 in AUPRC compared to DMSR without features.

**Table I.** AUROC and AUPRC for the three compared models. Blue and bold type indicates best results.

| Model | AUROC | AUPRC |
| --- | --- | --- |
| REDIRECTION | 0.8700 | 0.8300 |
| DMSR-WF | 0.9028<br>± 0.0079 | 0.9177<br>± 0.0041 |
| DMSR | **0.9148**<br>± **0.0070** | **0.9219**<br>± **0.0036** |

The progress made is noteworthy, given the significantly good results of the REDIRECTION model. It demonstrates that node features are essential in GNNs and improve the outcomes. However, to ensure a more equitable comparison, each model should undergo hyperparameter tuning to ensure that they are operating at their best, but time constraints prevented this from being done. Performing hyperparameter tuning could potentially result in an even greater improvement margin. Since the same set of hyperparameters were used for training models on datasets with varying levels of information, the performance of more complex datasets may have been hindered. This is likely due to the fact that the hyperparameters were heuristically set for the dataset with the least information.

Another way to evaluate and compare the models is by analyzing the predicted value distribution for the RepoDB test and random edges. A desirable distribution would be one in which all random edges have a predicted value of 0, while all RepoDB edges have a predicted value of 1. As shown in Fig. 3 and Fig. 4, both models perform well, with the distribution of prediction scores for random edges skewed towards 0 and the distribution for RepoDB edges skewed towards 1. This indicates that both models are capable of distinguishing between positive and negative drug repurposing hypotheses.

**Fig. 3.**
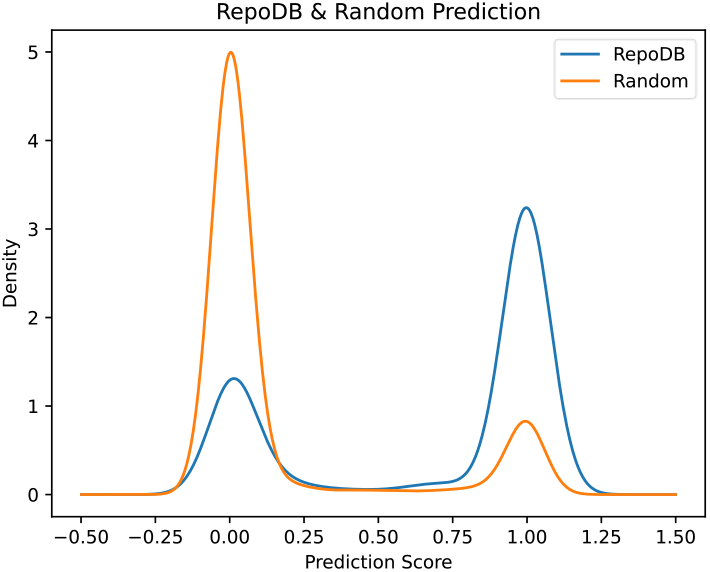
Distribution of REDIRECTION scores. RepoDB validating set in blue and a randomly generated set of links in orange.

**Fig. 4.**
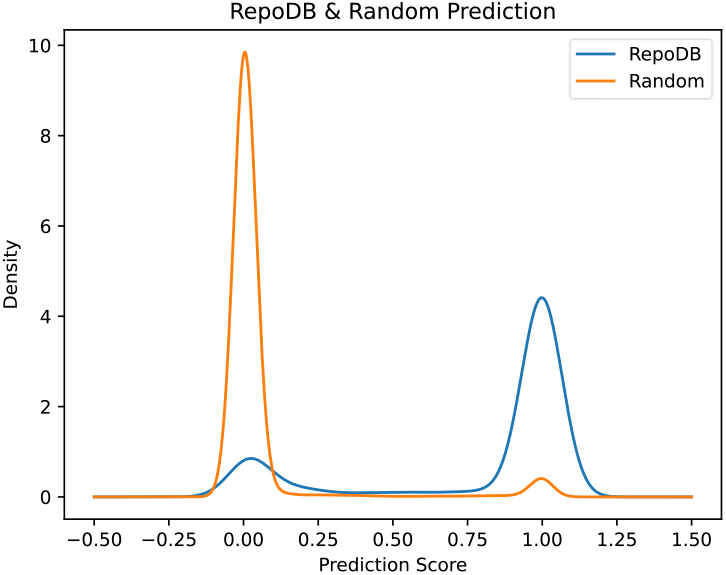
Distribution of DMSR scores. RepoDB validating set in blue and a randomly generated set of links in orange.

When compared to REDIRECTION, the results of the presented model in this work show a notable improvement, as can be seen by comparing Fig. 3 and Fig. 4 (note that the scales for the y axis are different). The main improvement lies in DMSR’;s ability to correctly identify random cases, resulting in a significant reduction of false positives and an increase in precision. On the other hand, the identification of positive cases has had an improvement as well.

TABLE II includes top 15 drug repurposing hypotheses retrieved by DMSR. All of them have a prediction score of 1. These edges were found using just the information of diseases, symptoms, drugs and their relationships, further analysis of these cases is necessary to asses their potential, but it’;s worth noting that the model was able to predict these potential relationships within a matter of seconds during inference.

**Table II.** Top 15 drug repurposing hypotheses retrieved by the model.

| Disease UMLS CUI | Disease Name | Drug ChEMBL ID | Drug Name |
| --- | --- | --- | --- |
| C0751137 | Craniofacial Pain | CHEMBL1490 | Trihexyphenidyl |
| C0086237 | Epilepsy, Cryptogenic | CHEMBL641 | Atomoxetine |
| C0270824 | Visual epilepsy | CHEMBL3833361 | Promethazine Teoclate |
| C3151568 | Nephrotic Syndrome, Type 4 | CHEMBL121 | Rosiglitazone |
| C0086237 | Epilepsy, Cryptogenic | CHEMBL640 | Procainamide |
| C0270824 | Visual epilepsy | CHEMBL3833362 | Zinc Oleate |
| C0270824 | Visual epilepsy | CHEMBL3833364 | Hydrargaphen |
| C0270824 | Visual epilepsy | CHEMBL3833368 | Rucaparib Camsylate |
| C0270824 | Visual epilepsy | CHEMBL3833369 | Fish Oil |
| C0270824 | Visual epilepsy | CHEMBL3833373 | Avelumab |
| C0270824 | Visual epilepsy | CHEMBL3833381 | Merbromin |
| C0270824 | Visual epilepsy | CHEMBL3833382 | Prajmalium |
| C0270824 | Visual epilepsy | CHEMBL3833383 | Gallium Dotatate GA-68 |
| C0270824 | Visual epilepsy | CHEMBL3833311 | Dextromoramide Tartrate |
| C0270824 | Visual epilepsy | CHEMBL3833388 | Prajmalium Bitartrate |

## V. Conclusions and future lines

Embedding the two-dimensional molecular structure of drugs using GNNs is an effective method to encapsulate useful information. Through the application of GNNs, the structure can be transformed into a representative vector that can serve as a feature for drug nodes in the broader task of generating hypotheses for drug repurposing.

Evidence has demonstrated that incorporating node features supports the model in the training phase, enabling it to learn the latent information necessary to address the task of generating hypotheses for drug repurposing. As a result, including these features in the DMSR model has led to improved outcomes compared to both the previous REDIRECTION and the enhanced model that did not include these features.

Regarding future directions, given the demonstrated significance of node features, the integration of such features for the phenotype node type appears to be the most critical area for advancement. Following this, expanding to a broader graph, similar to the one presented in Ayuso Muñoz [29], could represent a promising avenue for exploration. Additionally, hyperparameter optimization presents an intriguing possibility for further research. It is worth noting that DMSR remains a prototype, and future improvements and refinements will involve incorporating additional information of varying types into the input graph, as well as augmenting the architecture with other expanded details.

Nonetheless, it should be noted that DMSR’s predictions cannot be considered as a definitive truth and require experts and experimental validation. It is crucial to understand that DMSR’s should not be viewed as a substitute for medical prescription systems. Rather, it serves as a tool to suggest potential drug repurposing pairs that may be worth further investigation.

## Acknowledgment

The work is a result of the project “Data-driven drug repositioning applying graph neural networks (3DR-GNN)“, that is being developed under grant “PID2021-122659OB-I00“ from the Spanish Ministerio de Ciencia e Innovación. This work has been supported by project MadridDataSpace4Pandemics, funded by Comunidad de Madrid (Consejería de Educación, Universidades, Ciencia y Portavocía) with FEDER funds as part of the response from the European Union to COVID-19 pandemia. Andrea Álvarez-Pé z’ was granted by Universidad Politécnica de Madrid and Banco Santander for a predoctoral Programa Propio grant. Belén Otero-Ca a c ’ work is supported by “Formation de Personal Investigador“ grant (FPI PRE2019-090912) as part of the project “DISNET (Creation and analysis of disease networks for drug repurposing from heterogeneous data sources applied to rare diseases)“ (RTI2018-094576-A-I00) from the Spanish Ministerio de Ciencia, Innovation y Universidades.

## Footnotes

1 https://medal.ctb.upm.es/internal/gitlab/disnet/gnns/cbms2023-gnns

